# Standardization of inducer-activated broad host range expression modules: Debugging and refactoring an alkane-responsive AlkS/*P_alkB_* device

**DOI:** 10.1101/2020.12.26.424440

**Authors:** Alejandro Arce-Rodríguez, Ilaria Benedetti, Rafael Silva-Rocha, Víctor de Lorenzo

**Affiliations:** Systems Biology Department, Centro Nacional de Biotecnología-CSIC, Campus de Cantoblanco, Madrid 28049, Spain

**Keywords:** SEVA, AlkS, *Pseudomonas*, Crc, Hfq, cytometry, noise

## Abstract

Although inducible heterologous expression systems have been available since the birth of recombinant DNA technology, the diversity of devices and genetic architectures of the corresponding vectors have often resulted in a lack of reproducibility and interoperability. In an effort to increase predictability of expression of genes of interest in a variety of possible bacterial hosts we propose a composition standard for debugging and reassembling all regulatory parts that participate in the performance of such devices. As a case study we address the *n*-octane and dicyclopropyl ketone (DCPK)-inducible *P*_*alkB*_ promoter of the alkane biodegradation pOCT plasmid of *Pseudomonas putida*. The standardized expression module consisted of an edited *alkS* regulatory gene that is divergently expressed and separated of *P*_*alkB*_ by a synthetic DNA buffer sequence. The native DNA sequence of the structural *alkS* gene was modified to alleviate the catabolite repression exerted by some carbon and nitrogen sources through the Crc/Hfq complex of some hosts. The *P*_*alkB*_ promoter along with the *alkS* variants were then formatted as SEVA (Standard European Vector Architecture) cargoes and their activity parameters in *P. putida* determined with GFP and luminiscent reporters. The thereby refactored system showed improvements in various features desirable in conditional expression modules: inducibility, capacity, noise reduction and on/off ratio. When applied to other promoter/regulator pairs, the compositional standard thereby implemented in the AlkS/*P*_*alkB*_ module will enable more complex genetic programming in non-model bacteria.

## INTRODUCTION

Heterologous expression of genes in hosts (e.g. bacteria) different from their native origin and triggered by an external inducer is one of the bases of modern Biotechnology (1). A large number of genetic devices to this end have been developed over the years following the pioneering use of the IPTG-inducible *lac* promoter for expression in *E. coli* (2). Typical modules involve one promoter that is activated or repressed by a cognate transcription factor (TF; either an activator or a repressor), which binds the target sequence or changes its activity in a fashion dependent on addition to the medium of a chemical inducer. This basic scheme is the template for a large number of popular expression vectors based on a suite of regulator/promoter pairs (1, 3, 4). While they have been useful when the issue was to express one or few genes at time in one host, the onset of Synthetic Biology in recent years has multiplied the need of multiple, regulatable promoters endowed with specific parameters and as independent as possible of the physiological state of the host (4). One step in that direction was the creation in 2013 of the so-called Standard European Vector Architecture (5), which compiles a large number of standardized antibiotic marker genes, broad host range origins or replication and functional cargoes, aimed at simplifying genetic programming of a wide variety of bacteria of industrial and environmental interest (6). One type of such cargoes includes expression modules. While the boundaries of the corresponding DNA segments within the plasmid vector or transposon vector frame are well defined in the SEVA format, the organization of the regulatory elements inside the cargo have not been standardized yet. In this work we propose a specific arrangement for such inducible modules that attempts to preserve the inducibility of the TF/promoter pairs usable for heterologous expression while de-compressing and simplifying the native regulation of the cognate systems.

As a case study for such standardization effort we have chosen the regulatory node that controls expression of the *alk* genes for biodegradation of octane borne by the so-called OCT plasmid of soil bacterium *Pseudomonas putida* GPo1 (7). In its native arrangement, two gene clusters are involved in the process. *alkST* encodes the *n-*octane-inducible transcriptional regulator of the pathway (AlkS) and AlkT (a component of alkane hydroxylase). The second *alkBFGHJKL* cluster determines the rest of the activities, which are expressed from the upstream AlkS-dependent promoter *P*_*alkB*_ (8, 9). Once excised from its native context and assembled adjacent to each other in a single DNA segment, the *alkS*/*P*_*alkB*_ pair has been used to develop a number of biosensors for alkanes as well as heterologous expression vectors (10–13). The last is facilitated by the use of the gratuitous, soluble inducer dicyclopropyl ketone (DCPK). However, this simple rearrangement of functional segments with *alkS* and *P*_*alkB*_ does not eliminate the regulatory complexity embodied in them. *alkS* is transcribed through two promoters, *P*_*alkS1*_ and *P*_*alkS2*_, which are negatively and positively regulated, respectively by AlkS. In addition, translation of AlkS is subject to the post-transcriptional control of the Crc/Hfq complex, which enters a catabolite repression signal in the system (14). Finally, the activity of AlkS seems to be influenced also by the cytochrome terminal oxidase Cyo (15). While such a regulatory density allows the extant system to compute many physiological signals other than the mere presence of pathway substrates, such intricacy is also a nuisance for the predictability of the gene expression module. In the work presented below we have constructed a refactored AlkS/*P*_*alkB*_ device in which any known regulatory control—other than induction by DCPK— has been erased and replaced by non-regulatory DNA sequences, following a defined composition standard. As shown below, the resulting inducible expression module displays an improved performance as compared to its counterparts with the wild-type sequences. On this basis, we advocate the general application of the compositional standard used to assemble this device for increasing the reproducibility and interoperability of a large number of other devices made with regulatory parts mined from the genomes of environmental bacteria.

## MATERIALS AND METHODS

### Strains, plasmids, and growth conditions

The bacteria and the plasmids used in this work are listed in Table 1. All *Pseudomonas putida* strains were derived from the reference variant KT2440 or its rifampicin resistant derivative KT2442. The *E. coli* strains CC118 and DH5*α* were used as hosts for maintenance of plasmids. Unless indicated otherwise, cells were grown at 30°C (*P. putida*) or 37°C (*E. coli*) in rich LB medium (16) amended, where necessary, with 100 µg/ml ampicillin (Ap) or 50 µg/ml streptomycin (Sm) to retain plasmids. For solid media preparation, LB medium was supplemented with 1.5% (w/v) Bacto Agar (Pronadisa). Where indicated, the expression of *P*_*alkB*_ promoter was induced by the addition of dicyclopropylketone (DCPK) in solid and liquid media at the concentrations indicated.

**Table 1.**
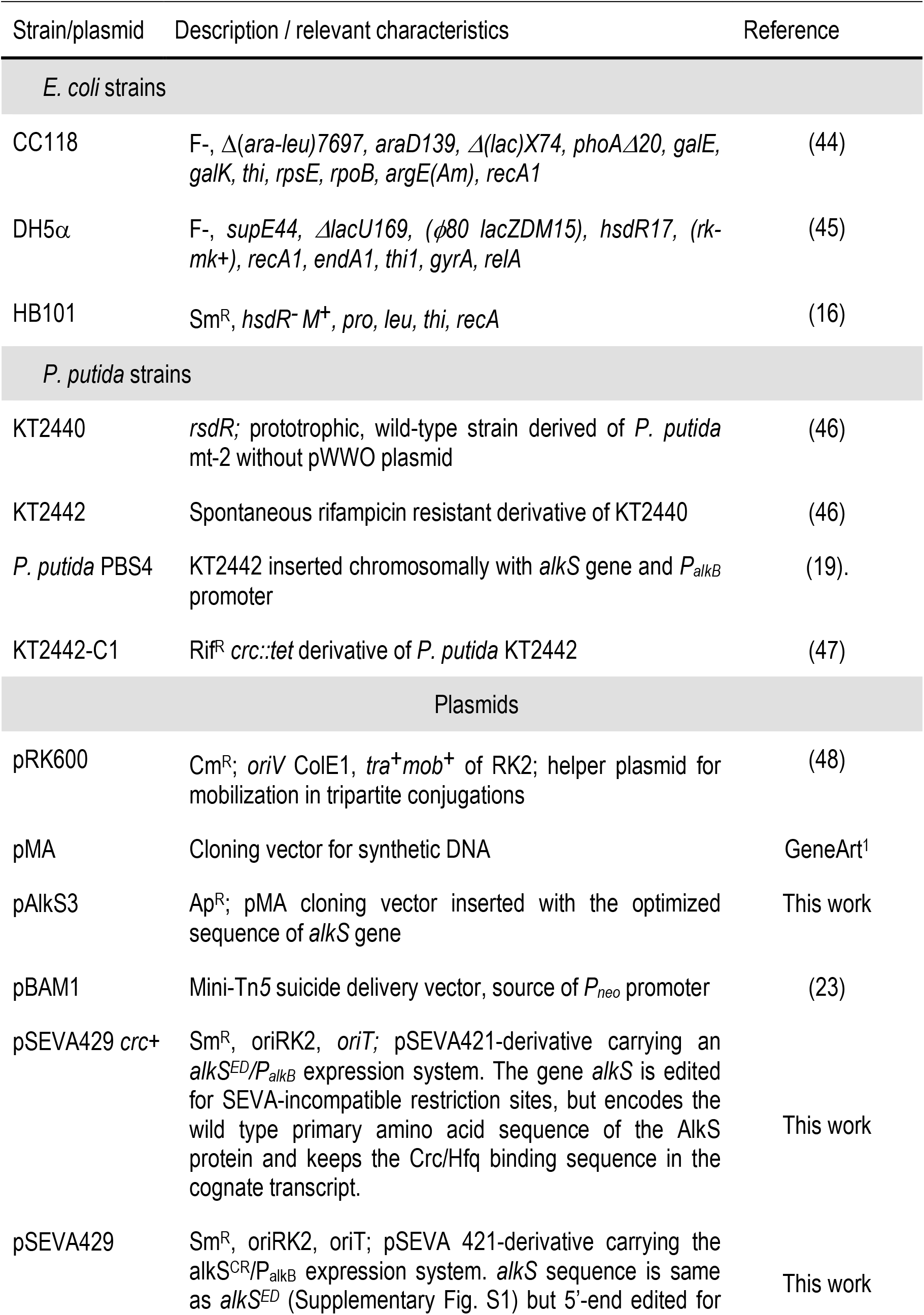
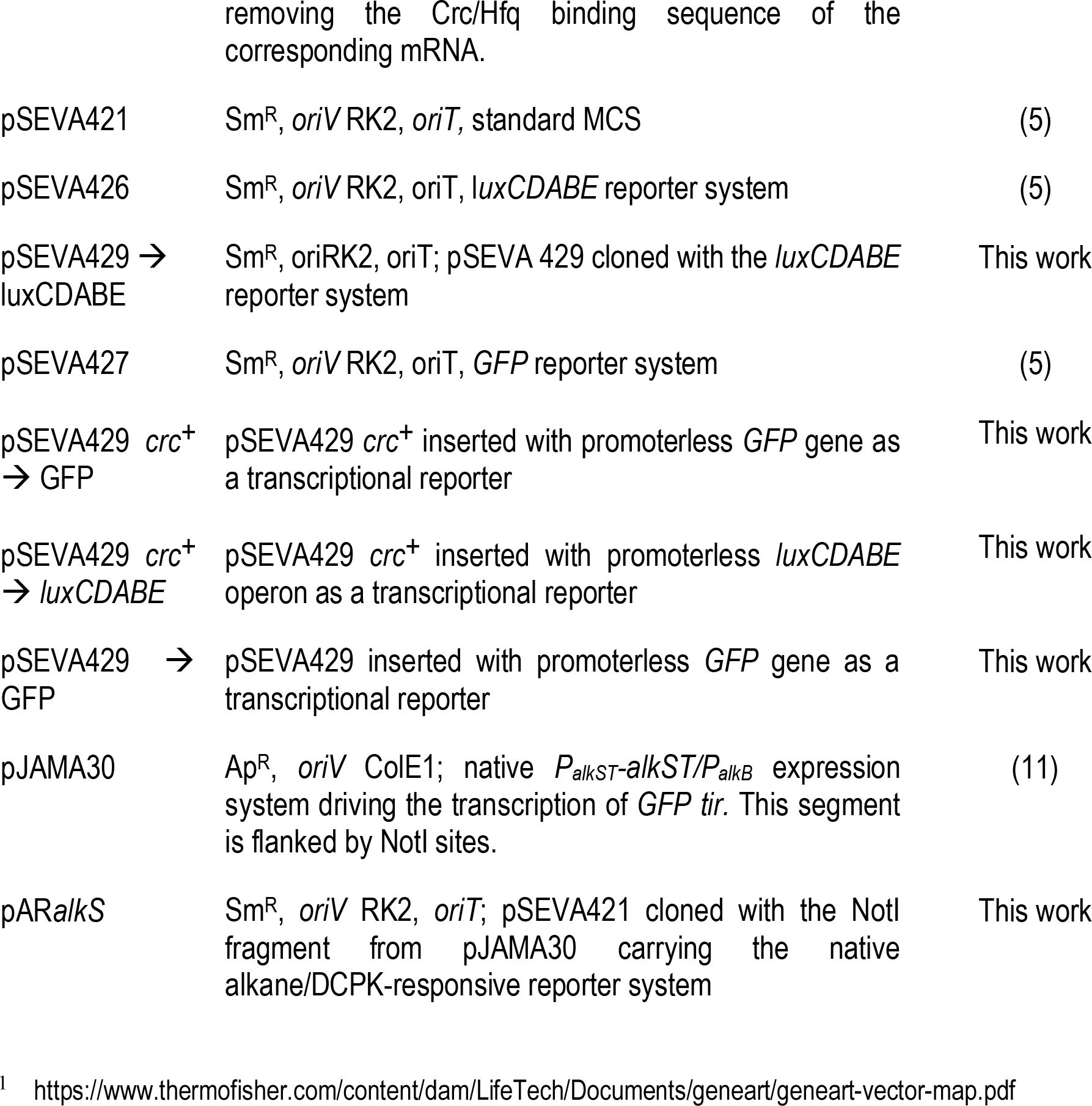
Strains and plasmids used in this work.

### Recombinant DNA techniques

General methods for DNA manipulation were performed with standard protocols described elsewhere (16). The amplification of DNA fragments by polymerase chain reaction (PCR) was implemented in 50 µl reactions containing approximately 100 ng of genomic DNA or 10 ng of plasmid as template, 0.25 mM dNTPs, 25 pmol of each primer and 1 U of GoTaq DNA polymerase (Promega). Reactions were run by an initial denaturalization (5 min, 94°C) followed by 30 cycles of denaturalization (1 min, 94°C), annealing (1 min, 58°-64°C), extension (1-3 min at 72°C) and final extension (10 min, 72°C). PCR products were purified with the NucleoSpin® Gel and PCR Clean-up kit (Macherey-Nagel) and, when required, digested with restriction enzymes purchased from New England Biolabs. Plasmid DNA was isolated by means of the Wizard® Plus SV Minipreps DNA Purification system (Promega). *E. coli* cells were transformed with plasmids with the CaCl_2_ method (16). In the case of *P. putida*, plasmids were incorporated by conjugative triparental mating using the *E. coli* HB101 (pRK600) as helper strain (17) or by electroporation of cells previously washed and concentrated with 300 mM sucrose as in Choi *et al*. (18).

### Construction of the *P*_*alkB*_/AlkS expression modules compatible with the Standard European Vector Architecture (SEVA)

In order to create an expression system that could fit the SEVA plasmid platform (5, 6), the gene *alkS* from *Pseudomonas putida* GPo1 was edited to remove incompatible restriction sites (*alkS*^*ED*^). The modified sequence (Supplementary Fig. S1) was fully synthesized by GeneArt/Thermofisher (Waltham, Massachusetts) and delivered as an insert in the pMA vector that was called pAlkS3. Both the AlkS regulatory protein and the *P*_*alkB*_ promoter were then arrayed with the other DNA segments indicated in Fig. 1. The *P*_*alkB*_ promoter was PCR-amplified with primers aaPalkB1-F (5’**AGCGGATAACAATTTCACACAGGA**CGTGTTTTTCCAGCAGACGAC3’) and aaPalkB1-R (5’ATGA<u>CCTAGG</u>CTCTCGACATCTTAAACCTGAGC3’), using as template genomic DNA from *P. putida* PBS4 (19). The *P*_*neo*_ promoter was also amplified by PCR with oligonucleotides aaPKm-NcoI1-F (5’TAGAA<u>CCATGG</u>TTTTTCCTCCTTATAAAG3’) and aaM13-R24-rev (5’ TCCTGTGTGAAATTGTTATCCGCT 3’) from pBAM1 (Table 1). The sequence of primer aaM13-R24-rev is complementary to the 5’ end of primer aaPalkB1-F (bold characters), allowing the assembly of *P*_*neo*_ with *P*_*alkB*_ by SOEing PCR (20). Insertion of the resulting product into the NcoI/AvrII restriction sites of a pSEVA frame gave rise to expression vector pSEVA429 *crc*^*+*^ (Supplementary Fig. S2)

**Figure 1.**
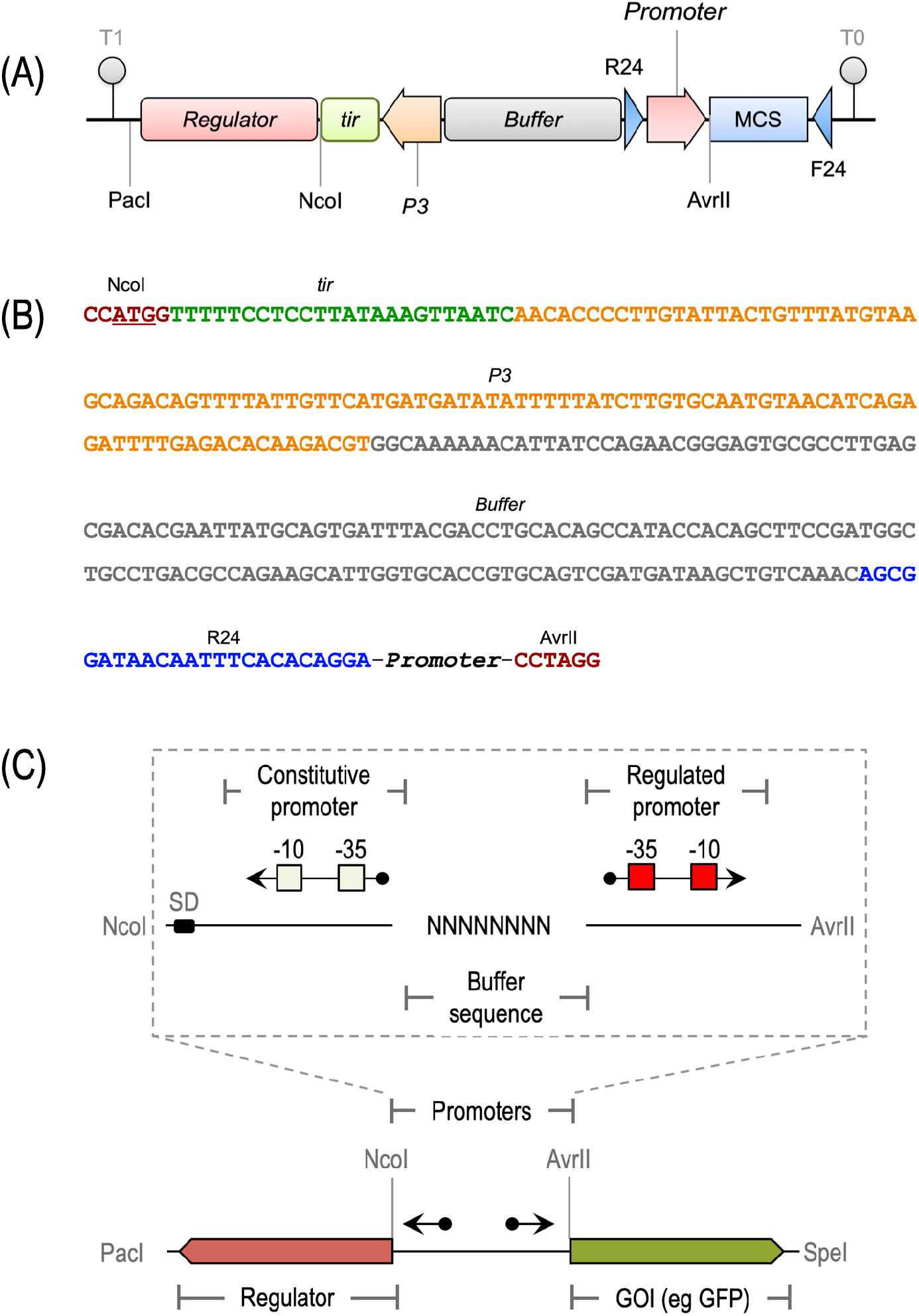
Organization of standardized inducible expression modules. (A) Arrangement of DNA portion, boundaries and their roles as the frame for inserting genes for inducer-responsive transcriptional regulators and cognate target promoters. (B) Blowup of the standardized DNA sequence that acts and the scaffold for the rest of the functional parts. (C) Configuration of reporter plasmids (with GFP or any other gene of interest, GoI) used in this work. Note constant and variable segments.

### Removal of the Crc binding site in the *alkS* gene

The Crc binding site in the 5’ end of the *alkS* gene in pSEVA429 *crc*^*+*^ was -modified with PCR-based site-directed mutagenesis. Briefly, the first 570 bp of *alkS* were amplified with the oligonucleotides 5-alkSmut-NcoI (5’GCGC<u>CCATGG</u>GCATGAA**G**AT**C**AA**G**AT**C**AT**C**AACAATGAT TTCCCGGTTGCCAAGATCG3’) and 3-alkSmut-XhoI (5’AGCGCCTGCAAGTTTAAGCC3’) using the pAlkS3 as template. The forward primer contains the recognition sequence of NcoI enzyme (underlined) and also 6 single-nucleotide mutations introduced deliberately to eliminate the Crc site from *alkS* (in bold characters). The PCR fragment was then restricted with the enzymes NcoI and XhoI (the last coded in between the *alkS* sequence) and the resulting 406 bp fragment was recloned into the same sites of pSEVA429 *crc*^*+*^. In order to follow the SEVA nomenclature, this final expression vector was named pSEVA429 (i.e. Sm resistance, RK2 origin of replication and AlkS/*P*_*alkB*_ expression cargo; Supplementary Fig. S2)

### Parameterization of the AlkS/*P*_*alkB*_ module

For generating constructs that report transcriptional activity as a fluorescent emission, the *GFP tir* gene of pSEVA427 was cloned into plasmids pSEVA429 *crc*^*+*^ and pSEVA429 as a HindIII/SpeI insert to generate plasmids pSEVA429 *crc*^*+*^→ GFP and pSEVA429 → GFP respectively. As a control, we excised the NotI fragment from pJAMA30 containing the original *alkST* genes of pOCT transcribed by their own *P*_*alkST*_ promoter and, in divergent orientation, the *P*_*alkB*_ driving the expression of the *GFP tir* gene (11; Supplementary Fig. S3). This ∼6.7 kb fragment was cloned into the NotI site of pSEVA421 to generate the control vector pAR*alkS*. The three *GFP tir* reporter vectors described above, were transferred into wildtype *P. putida* KT2440 cells and into the *crc::tet* derivative of *P. putida* KT2442. Plasmid-bearing strains were then grown at 30°C until mid-exponential phase, the cultures added with 0.05% v/v DCPK and fluorescent emission for the next 3 hours followed as described in detail in (21) with a Gallios (Perkin Elmer) flow cytometer. GFP was excited at 488 nm, and the fluorescence signal was recovered with a 525/40 BP filter. The resulting data was processed using FlowJo v. 9.6.2 software (FlowJo LLC, Ashland, OR, USA). For monitoring activity of the AlkS/*P*_*alkB*_ module at a population level, plasmids pSEVA429 *crc*^*+*^ and pSEVA429 were inserted with the promoterless luminescent reporter *luxCDABE* operon excised from pSEVA426 as a HindIII/SpeI fragment. This originated plasmids pSEVA429 *crc*^*+*^→ *luxCDABE* and pSEVA429 → *luxCDABE* respectively. As with the GFP counterparts before, these plasmids were passed to wild type and *crc*-minus *P. putida* hosts. For measuring light emission under various DCPK concentrations, cells grown overnight in LB were diluted in same medium, placed in a Microtest™ 96-well assay plate (BD Falcon), regrown to mid-exponential phase, added with the inducer and luminescence recorded after 4 h.

## RESULTS AND DISCUSSION

### A compositional standard for engineering inducer-dependent heterologous gene expression

Fig. 1A sketches the organization of the inducible expression module proposed in this work for SEVA cargoes (5) aimed at heterologous expression of genes of interest in a variety of bacterial hosts. First, the standard asks for constitutive expression of the gene(s) encoding the effector-responsive regulator. In their natural context, TFs are often subject to a degree of self-regulation, either positive or negative (22) that enters a layer of unnecessary complexity that the arrangement shown in Fig. 1A mitigates if not entirely eliminates. The standardized sequence that connects the various functional parts of the device (Fig. 1B) includes the following segments. The source of transcription of the signal-responsive TF gene is the 106 bp minimal promoter P3/*P*_*neo*_ that drives expression of the kanamycin resistance gene of pBAM1 (23). Following this promoter, default translation efficiency is also fixed by means of a 24 bp translation initiation region (TIR) retrieved from the GFP variant borne by pGreenTIR plasmid (24). This is an unusual ribosome binding sequence known to act as a translational enhancer that is expected to curb expression noise that could originate in a poor translation. The segment for constitutive expression/translation of the TF is followed upstream by a 150 bp neutral segment of DNA retrieved from the *lacI^q^-P*_*trc*_ expression system of plasmid pTrcA (25). This sequence has no known regulatory elements and it functions as a buffer region to ease mutually negative supercoiling that could stem from transcription of divergent promoters (26). As shown in Fig. 1A, the adjacent piece of DNA is the one that bears the promoter targeted by the inducible regulator and orientated opposite in respect to the sequences for expression of the TF gene. The specific DNA sequence of this promoter obviously changes from case to case but it should by default be accommodated within a segment of nor more than 100 pb. The 3’ of this promoter sequence is bound by an AvrII site, which links this segment to the start of the SEVA polylinker (5). Note that two sites at the boundary buffer sequence/promoter and at the end of the MCS have target sequences for oligonucleotides R24 and F24 (5) for easy amplification and diagnose of possible inserts. Once the gene encoding the inducer-responsive TF is placed in this arrangement as a NcoI (overlapping the leading ATG)-PacI DNA fragment, the whole expression module becomes inserted in the SEVA frame as a PacI-AvrII addition (Fig. 1C), shielded both upstream and downstream by transcriptional terminators contributed by the vector structure—and ready to be inserted with any gene of interest (GOI) cloned in the corresponding polylinker.

### Reshaping the *alkS* and *P*_*alkB*_ pair as an inducible expression device

As a case study of formatting a naturally-occurring inducible promoter into a standardized expression cargo we picked the regulatory elements that control transcription of the *alk* genes of the OCT plasmid of *Pseudomonas putida* GPo1 (7, 8). The choice was motivated by the exemplary regulatory density of the extant system, that includes transcriptional and post-transcriptional control layers checking expression of AlkS (Fig. 2A). This provided an archetypal case to inspect the impact of the simplified formatting explained above on the behaviour of the resulting expression module. To this end, we first edited the wild-type DNA sequence of the regulator to eliminate restriction sites incompatible with the SEVA rules while preserving the primary amino acid sequence. The resulting DNA segment was then produced as a 5’ → 3’ 2668 bp NcoI-PacI DNA fragment. The complete list of changes entered in this *alkS* variant (that we term *alkS*^*ED*^ for edited) is compiled in Supplementary Fig. S1. The *alkS*^*ED*^ variant was then coupled to the spacer shown in Fig. 1B, which was added with the wild-type 90 bp sequence of the target *P*_*alkB*_ promoter (see Materials and Methods). The resulting cargo was subsequently placed in the frame of plasmid pSEVA421 (6) as a PacI-AvrII insert, resulting in expression vector pSEVA429 *crc+* (Table 1; Supplementary Fig. S2). For parameterization of the activity of the thereby refactored expression device, the construct was added with the promoterless GFP *tir* gene of pSEVA427 (6) downstream of *P*_*alkB*_ (Materials and Methods) resulting in reporter plasmid pSEVA429 *crc+* → GFP. To have a control of the wild-type expression device endowed of all native regulatory parts present in plasmid pOCT we excised the ∼6.7 kb NotI fragment of plasmid pJAMA30 (11) containing *alkST* and a divergent *P*_*alkB*_ promoter upstream of a GFP reporter (Supplementary Fig. S3). This DNA was then inserted into the same plasmid frame of pSEVA421 used for the edited devices, thereby originating control plasmid pAR*alkS* (Table 1; note that the *alkS* variant in this case is the very original wild type *alkS*^*WT*^ as shown in Supplementary Fig. S1).

**Figure 2.**
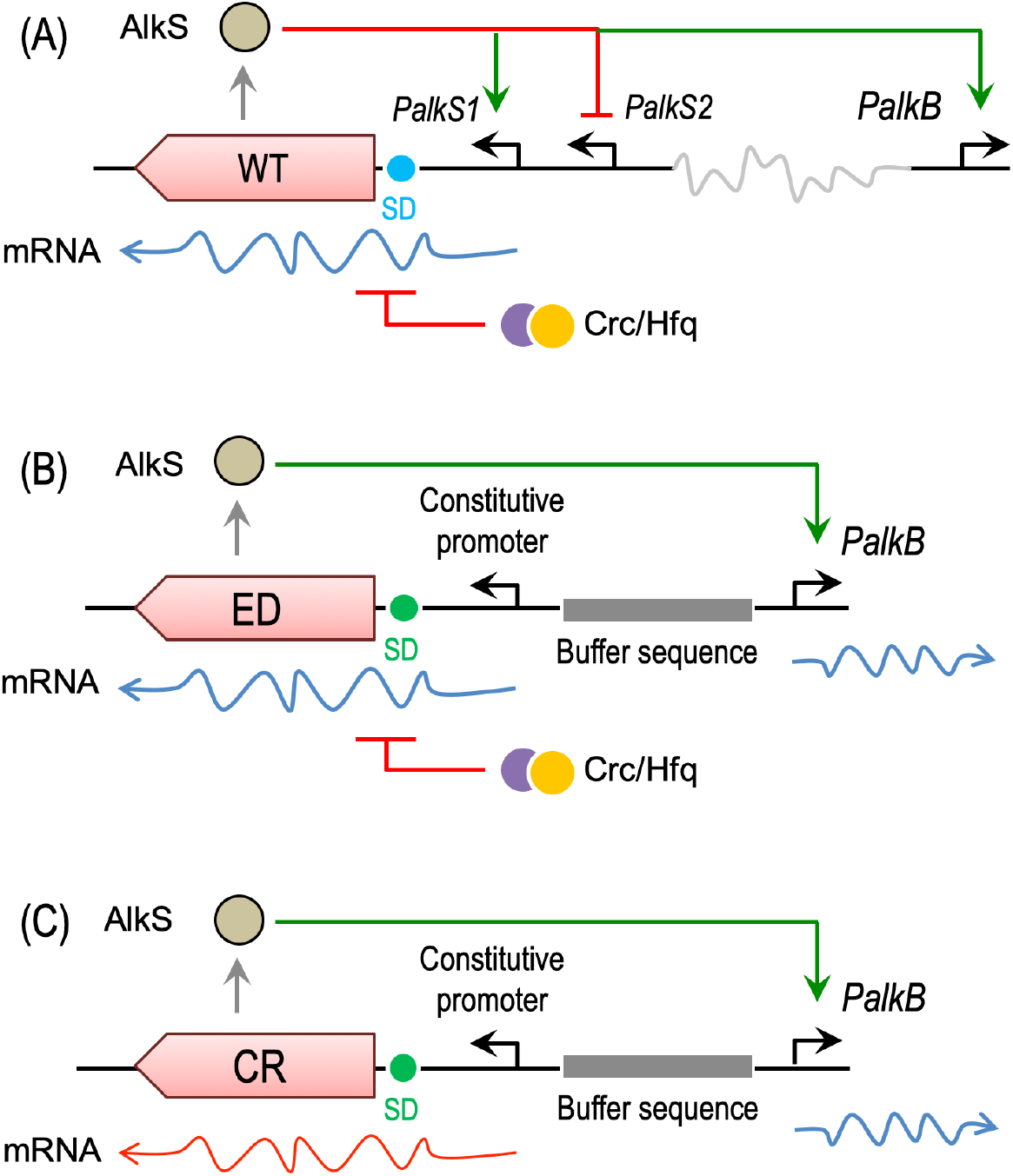
Functional segments of the naturally-occurring and standardized AlkS/*P*_*alkB*_ device. (A) Native organization of regulatory parts in the original context of pOCT plasmid. Note complex arrangement of transcriptional and post-transcriptional signals (e.g. inhibition of *alkS* mRNA translation by the Crc/Hfq complex) and dual effect of inducer-activated AlkS on self-promoters *P*_*alkS1*_ and *P*_*alkS2*_. (B) Constitution of the standardized AlkS *P*_*alkB*_ expression module. The wild-type DNA sequence of the regulator has been edited to remove restriction sites incompatible with the SEVA standard but keeping the same primary amino acid sequence (AlkS^ED^). *alkS* expression is now under the control of a heterologous SD sequence and a constitutive promoter (*P*_*neo*_, see text) and closer to target, divergent promoter *P*_*alkB*_ —albeit separated by the buffer sequence indicated in Fig. 1B. (C) AlkS/ *P*_*alkB*_ device bearing a regulator variant erased of its Crc binding site. As before, this change in the DNA of *alkS* keeps the primary amino acid sequence of the protein identical to the wild-type regulator.

Once equivalent constructs with *alkS*^*WT*^ (pAR*alkS*) and *alkS*^*ED*^ (pSEVA429 *crc*^*+*^ → GFP) were in hand, we were able to evaluate the effect of the standardization of the architecture of the regulatory node on transcriptional performance.

### The effect of formatting AlkS/*P*_*alkB*_ in the performance of the expression system

In a first series of experiments we compared the behaviour of the AlkS/*P*_*alkB*_ pair assembled with all the native regulatory system borne by the pOCT plasmid (Fig. 2A) and recreated in plasmid pAR*alkS* versus that of the same regulatory system arranged with the composition standard of Fig. 1 as implemented in pSEVA429 *crc*^*+*^ → GFP. Differences included a synthetic expression segment for transcription and translation of the regulator and an upstream buffer DNA sequence that was followed by the divergent AlkS target promoter *P*_*alkB*_ as sketched in Fig. 2B. Wild-type P. putida KT2440 was transformed with each of these two plasmids and transformants were grown in LB medium with Sm and fluorescent readout followed as explained in the Materials and Methods section. The cytometry results of these experiments are shown in Fig. 3. Inspection of the resulting graphs revealed some features of both the native and the formatted system that are worth to consider for handling of the expression device. First, formatted or not, the *P*_*alkB*_ has a noticeable non-induced basal transcription level that spontaneously increased with growth (Fig. 3A). Although this level is not too high (it remains within the same order of magnitude than the baseline expression), it has to be taken into account when expression of toxic proteins or coupling with other devices are pursued (27). A second aspect of the results shown in Fig. 3 is the noticeably stronger and faster induction of the standardized module of pSEVA429 *crc*^*+*^ → GFP as compared to the original regulatory node of pAR*alkS*. Finally, both devices displayed a growingly sharp unimodal expression pattern (28) along the induction period with little or no significant noise or cell-to-cell variation. While these results were good enough to accredit the correct functioning of the expression device following reassembly of its DNA parts, we wondered about the less predictable effects of other physiological inputs that operate on the AlkS/*P*_*alkB*_ in its native context.

**Figure 3.**
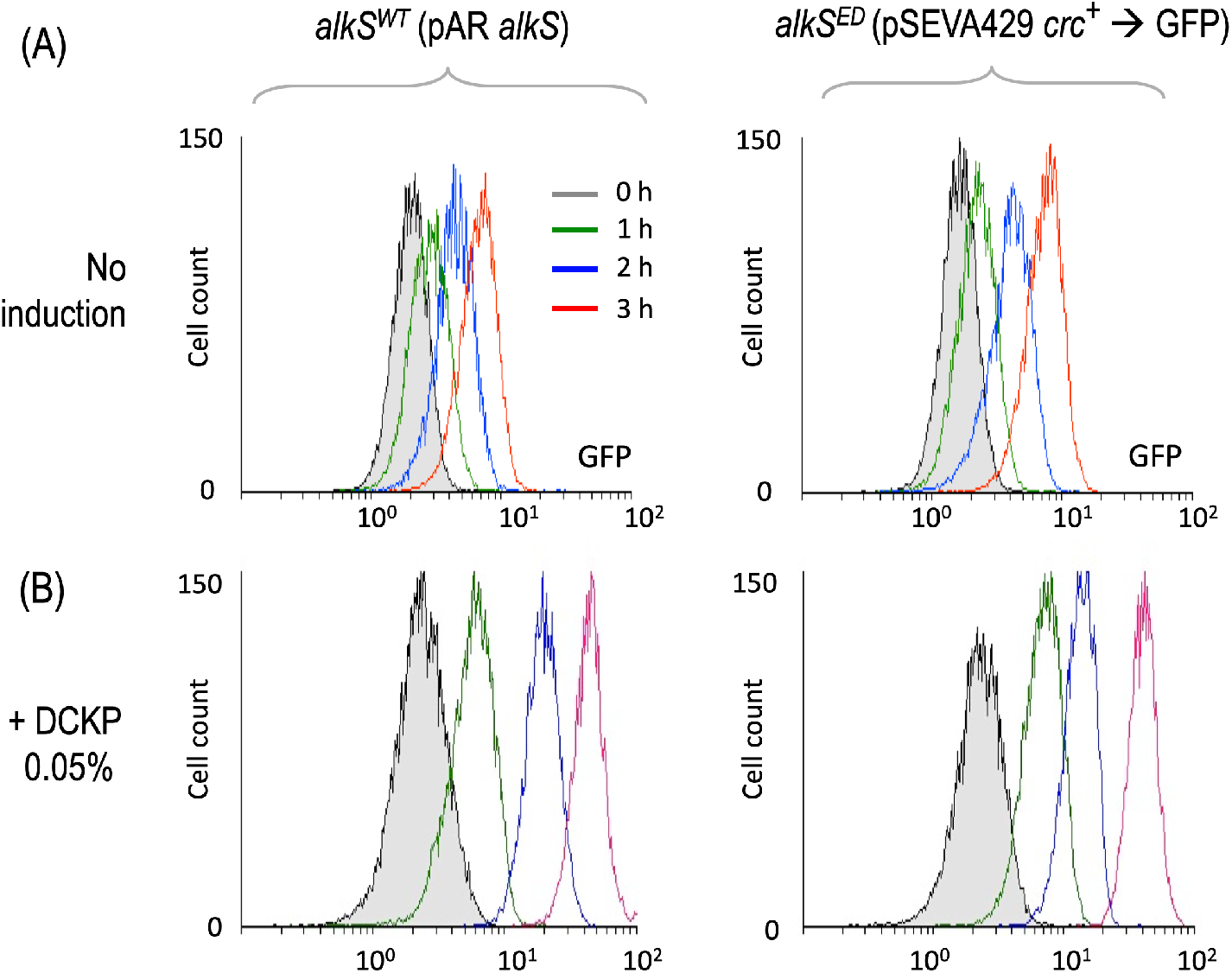
Transcriptional activity of AlkS/ *P*_*alkB*_ node before and after standardization as SEVA cargo. *P. putida* transformants with either pAR*alkS* (wild-type *alkS*) and pSEVA429 *crc*^*+*^ → GFP (edited *alkS* variant ED) were grown in LB until mid-exponential phase and added with 0.05% AlkS effector DCPK. Fluorescent emission was then measured in a cytometer for the next 4 hours as explained in the Materials and Methods section. (A) Control, no induction. (B) Cultures supplemented with inducer.

### Effect of Crc on the formatted and not-formatted AlkS/*P*_*alkB*_ device

Ideally, for engineering reliable genetic devices regulatory parts should deliver their function in a context-independent manner (29). Such a context includes not only genomic locations (30) and availability of resources (31), but also physiological signals (32, 33) that orchestrate the induction hierarchy. One of these is catabolite repression (34), which in the case of *P. putida* operates through a complex interplay between proteins Crc and Hfq with small RNAs to inhibit translation of mRNAs of target genes (14). AlkS happens to be subject to such post-transcriptional regulation when placed in *P. putida*, but not in *E. coli* (35). In order to calibrate the effect of such a control layer and whether it was kept or not in the standardized construct, we run the experiments shown in Fig. 4. To this end pAR*alkS* and pSEVA429 *crc*^*+*^ → GFP were placed in a *crc::tet* mutant of *P. putida* (Table 1) known to be blind to catabolite repression caused by many of the components of LB medium (36, 37). Although not as intense as reported in the native configuration an effect of lacking Crc became evident, specifically on the net accumulation of reporter signal in both constructs as well as an increase on the noise of basal promoter activity at longer growth periods. But otherwise, the overall behaviour of the standardized and non-standardized expression devices remained basically identical.

**Figure 4.**
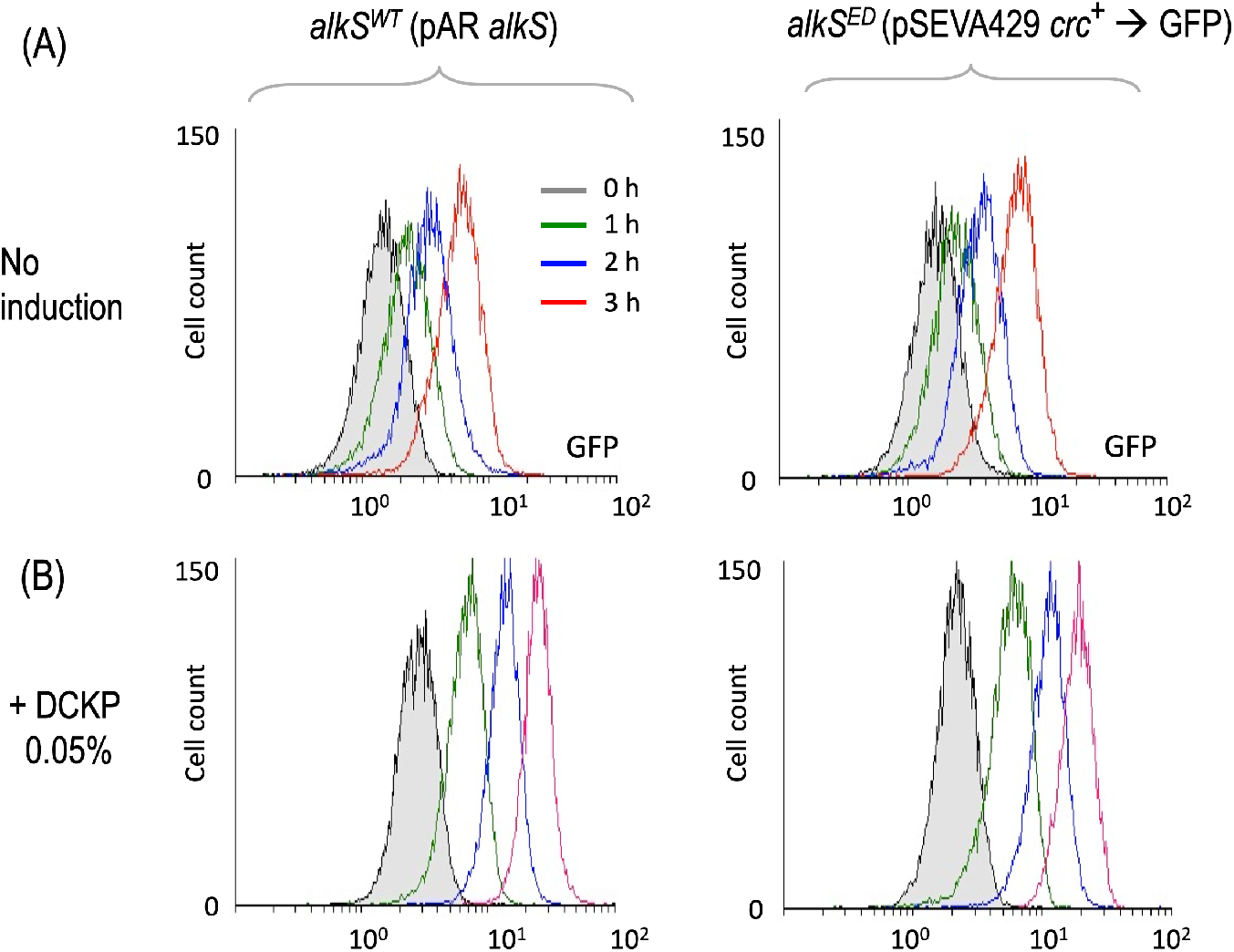
Effect of Crc on performance of the standardized AlkS/ *P*_*alkB*_ module. Plasmids pAR*alkS* (*alkS*^*WT*^) and pSEVA429 *crc*^*+*^ → GFP (*alkS*^*ED*^) were placed in a *crc::tet* mutant of *P. putida*, the transformants grown in LB and treated as indicated in the legend of Fig. 3. (A) Control, no induction. (B) Cultures supplemented with inducer 0.05 % DCPK.

The data above were welcome results, as refactoring of an existing regulatory node with a different architecture often results in devices that do worse than the naturally-occurring setup (38, 39). But the question still remained of whether we could erase altogether the effect of *crc* on the performance of the AlkS/*P*_*alkB*_ module, not by playing with the genetic background but by rewriting the DNA sequence of the regulator. To address this, an additional *alkS* derivative was synthesized in which the Hfq/Crc binding site of the corresponding mRNA 5’-AATAAAAATAATAAA-3’ that overlaps amino acid positions 4-9 of the protein was edited to 5’-GATCAAGATCATCAA-3’. As shown in Supplementary Fig. S1, these changes are predicted to abolish altogether the target site for Hfq/Crc, but they still keep the wild-type primary amino acid sequence of the encoded AlkS. As before, the resulting DNA (that bears the variant hereafter called *alkS*^*CR*^ for Crc-free) was formatted as a NcoI-PacI fragment and coupled to the spacer shown in Fig. 1B and the *P*_*alkB*_ promoter. The resulting cargo was subsequently placed in the frame of plasmid pSEVA421 as PacI-AvrII insert, resulting in expression vector pSEVA429 (Supplementary Fig. S2). For the sake of comparing its performance, a promoterless GFP gene identical to that of pAR*alkS* and pSEVA429 *crc*^*+*^ → GFP, was added to pSEVA429, thereby generating pSEVA429 → GFP. This *alkS*^*CR*^ -containing plasmid was then placed in *crc*^*+*^ and *crc*^*-*^ strains of *P. putida* and the readout of the fluorescent reporters followed in LB medium with or without DCPK induction as before. As shown in Fig. 5, removal of the *crc* site from the *alkS* sequence had only a moderate effect on the performance of the expression device, as the induction patterns were quite similar when pSEVA429 → GFP was placed in isogenic *P. putida* strains with or without the factor.

**Figure 5.**
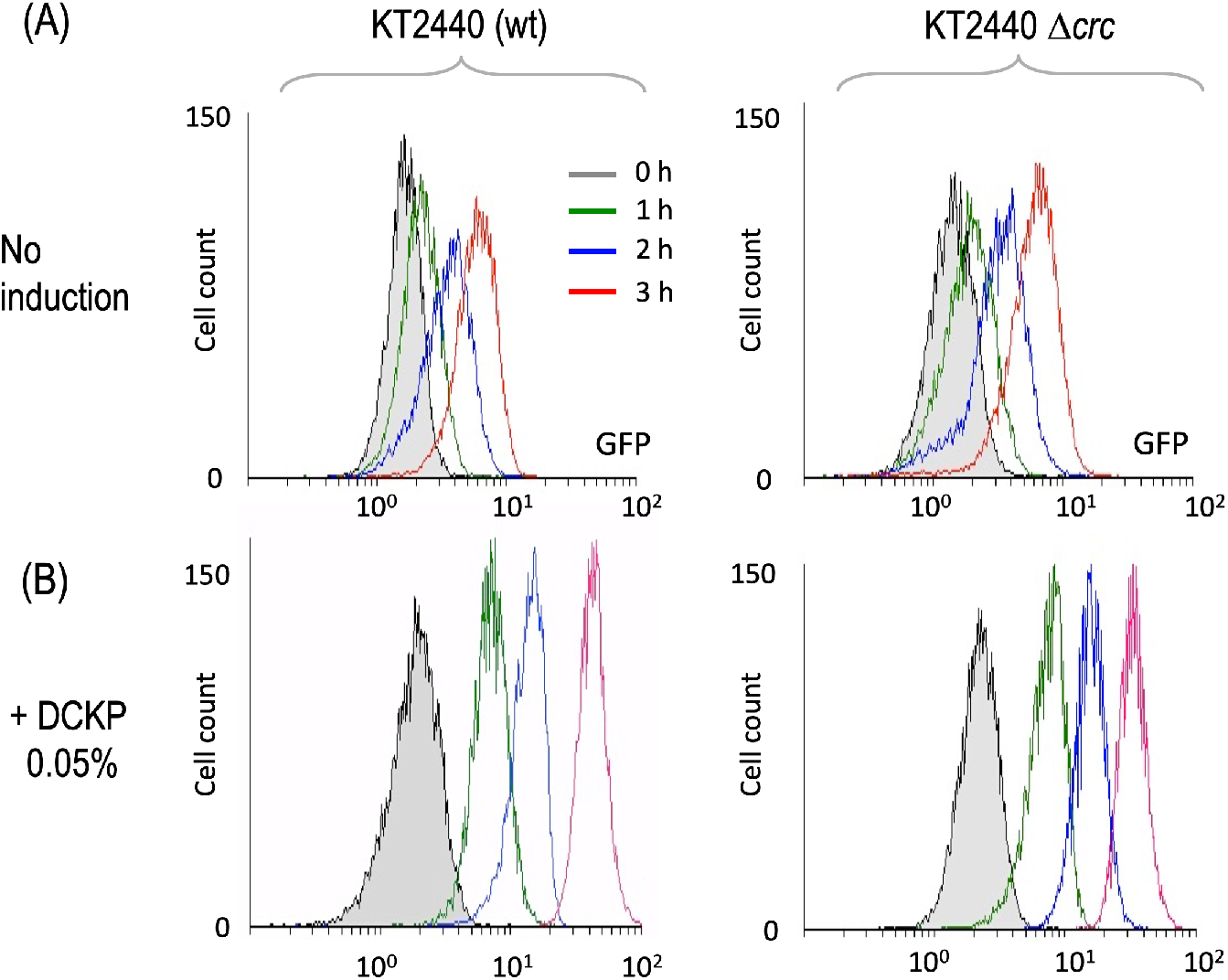
Behavior of an *alkS* gene variant erased of the Crc/Hfq binding site to its mRNA. Plasmid pSEVA429 → GFP (with the AlkS/*P*_*alkB*_ device bearing the regulator gene devoid of the Crc binding site in its mRNA) was transformed in *crc*^*+*^ and *crc*^*-*^ strains of *P. putida*. Transformants were grown in LB an treated as indicated in the legend of Fig. 3. (A) Control, no induction. (B) Cultures supplemented with inducer 0.05 % DCPK.

To gain some insight into this apparently minor influence of *crc* we constructed additional derivatives of pSEVA429 *crc*^*+*^ and pSEVA429 with a promoterless *luxCDABE* operon instead of GFP. This luminiscent reporter is way more sensitive than GFP and therefore a better proxy of transcriptional output from *P*_*alkB*_ at a population level. The new construct was placed as before in equivalent *crc*^*+*^ and *crc*^*-*^ strains and the cognate transformants grown in LB with different inducer concentrations. In this case (Fig. 6) both the overall weight of Crc on the system and the effect of removing its target site in *alkS* mRNA became clear. Working in an entirely Crc-free strain exposed to 0.05% DCPK equalized *P*_*alkB*_ activity at high levels regardless of whether the *alkS* variant of the expression vector had the *crc* site or not (Fig. 6C). In contrast, when placed in a wild-type strain, the construct with *alkS*^*CR*^ produced luminescence levels approximately twice as high those as the same with *alkS*^*ED*^. While this accredited an inhibitory role of Crc on AlkS, removal of the related sequence failed to make the AlkS/*P*_*alkB*_ altogether blind to catabolite repression—surely because this physiological control influences additional components of the system other than the transcriptional regulator

**Figure 6.**
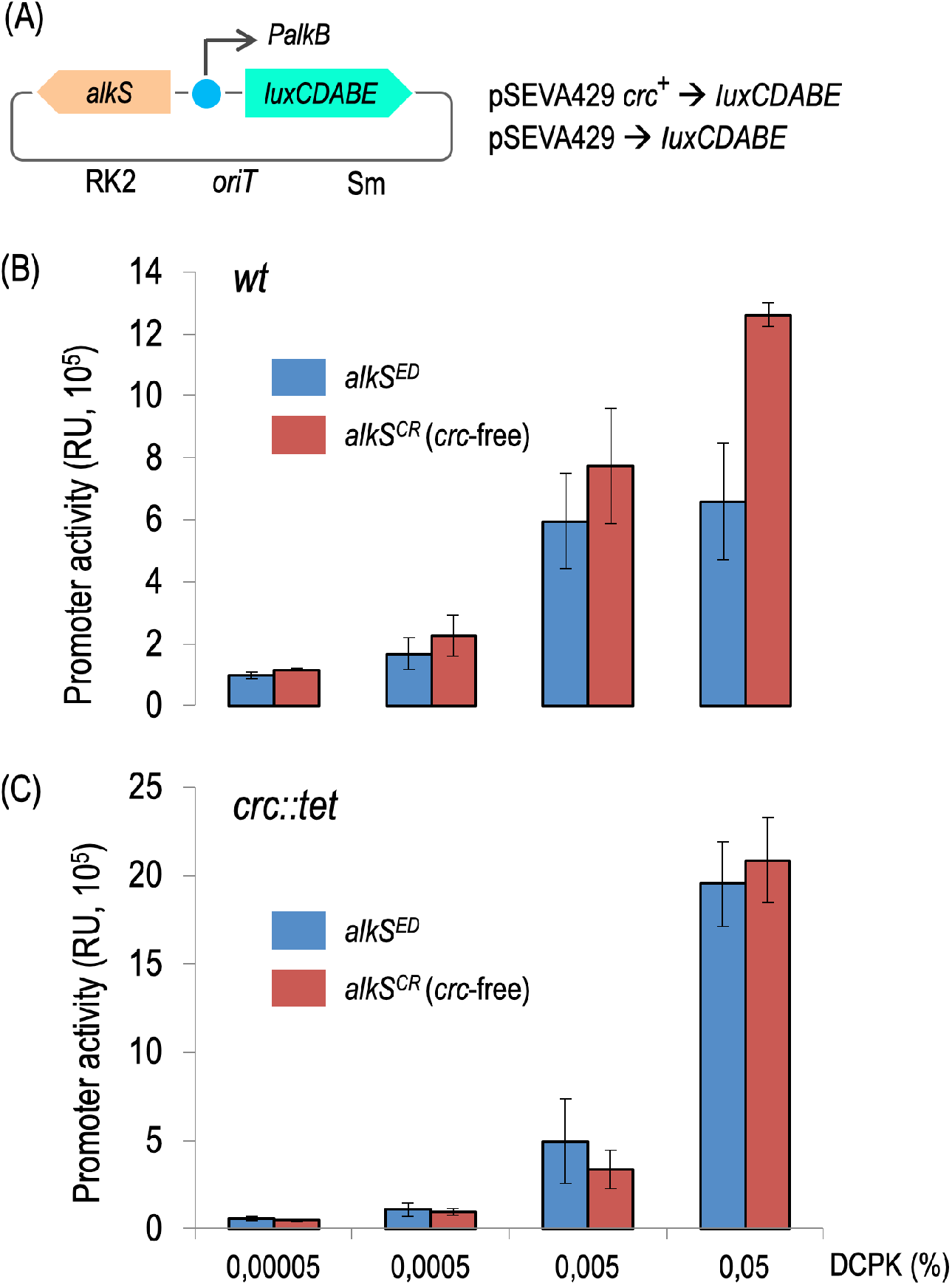
Influence of Crc at in *P. putida* populations bearing the AlkS/*P*_*alkB*_ module with the regulator gene with or without the Crc site. Plasmids pSEVA429 *crc*^*+*^ → *luxCDABE* (*alkS*^*ED*^, with Crc-binding site) and pSEVA429 → *luxCDABE* (*alkS*^*CR*^, without Crc site) were transformed in *crc*^*+*^ and *crc*^*-*^ strains of *P. putida*, grown in LB until mid-exponential phase and added with the DCPK concentrations indicated in each case. Luminiscent emission was then recorded after 4 h as a proxy of transcriptional activity as explained in the Materials and Methods section. (A) Sketch, not to scale, of functional segments in reporter plasmids. (B) Readout of reporter constructs borne by wild-type *P. putida*. (C) Same in *crc::tet* cells. Note different scales in Y axes.

## CONCLUSIONS

In this work we have used the inducer dependent and AlkS-mediated activation of the *P*_*alkB*_ promoter of the pOCT plasmid as an example of the roadmap that should be followed for reshaping a naturally-occurring regulatory node into an standardized device for engineering heterologous expression (40). As with any standard, there is a somewhat arbitrary but still reasonable and scientifically justifiable choice of a given composition rule (41). The one we propose in this work is summarized in Fig. 1 and explained in detail above. As is also the case of other standards, this particular choice will certainly limit flexibility but will foster interoperability, parameterization and comparative metrology (42, 43). The work above exemplifies how the same device, still after formatting, may go through successive, improved versions of the same functional DNA segments even if a prefixed arrangement is kept constant. In the cases examined above we can consider plasmids pAR*alkS* and pSEVA429 *crc*^*+*^ as beta versions of what we propose to be the *bona fide* standardized AlkS/*P*_*alkB*_ expression device apt for inclusion as a cargo in the SEVA collection: pSEVA429 (Supplementary Fig. S2). Note however that—as shown above—there is still room for improvement and it is likely that other versions will follow, an issue that is contemplated in the updated nomenclature of the SEVA collection (6). For instance, the system could be refactored to make it more digital (e.g. suppressing its considerable basal level; 27) and making it still more independent of catabolite repression. But, in the meantime, pSEVA429 is an altogether standardized off-the-shelf expression vector with a large number of benefits, including the possibility of comparing faithfully its performance with other expression modules that follow the same arrangement. We ultimately expect such standardization to ease engineering of complex systems and encourage other genetic tool developers to follow suit.

## SUPPLEMENTARY DATA

**Supplementary Figure S1**. Alignment of the wild-type *alkS* sequence with its edited variants *alkS*^*ED*^ and *alkS*^*CR*^.

**Supplementary Figure S2**. Organization of AlkS-based expression vectors

**Supplementary Figure S3**. Functional insert of plasmids pJAMA30 and pAR*alkS*

## Acknowledgments

Authors are indebted to Fernando Rojo and Renata Moreno for strains and valuable materials.

