## Supplementary Information for "Standardization of inducer-activated broad host range expression modules: Debugging and refactoring an alkane-responsive AlkS/*P_alkB_* device"

### SUPPLEMENTARY DATA

**Supplementary Figure S1.** Alignment of the wild-type *alkS* sequence with its edited variants *alkS*[ED] and *alkS*[CR].

```
CLUSTAL O(1.2.4) multiple sequence alignment

                           MetGlyMetLysIleLysIleIleAsnAsnAspPheProValAlaLysIleGlyLeuAsp
native_alkS                ------atgaaAATAAAAATAATAAAtaatgatttcccggttgccaagatcgggctggat
optimized_alkS-ED          atgggcatgaaAATAAAAATAATAAAtaatgatttcccggttgccaagatcgggctggat
optimized_alkS-CR          atgggcatgaaGatCaaGatCatCaaCaatgatttcccggttgccaagatcgggctggat
                                 ***** ** ** ** ** ** *********************************
                                        Crc binding

                           ArgIleThrThrLeuValSerAlaLysValHisAsnCysIleTyrArgProArgLeuSer
native_alkS                cgaattacgactctagtaagtgccaaagtgcataactgcatatatcggcccagattgagt
optimized_alkS-ED          cgaattacgactctagtaagtgccaaagtgcataactgcatatatcggcccagattgagt
optimized_alkS-CR          cgaattacgactctagtaagtgccaaagtgcataactgcatatatcggcccagattgagt
                           ************************************************************

                           IleAlaAspGlyThrAlaProArgValCysLeuTyrArgAlaProProGlyTyrGlyLys
native_alkS                atcgcggatggaactgcacccagagtatgcctttatagagccccgcctggatatggaaaa
optimized_alkS-ED          atcgcggatggaactgcacccagagtatgcctttatagagccccgcctggatatggaaaa
optimized_alkS-CR          atcgcggatggaactgcacccagagtatgcctttatagagccccgcctggatatggaaaa
                           ************************************************************

                           ThrValAlaLeuAlaPheGluTrpLeuArgHisArgThrThrGlyArgProAlaValTrp
native_alkS                accgtggctcttgcgttcgagtggctacgccacagaacaaccggacgtcctgcagtgtgg
optimized_alkS-ED          accgtggctcttgcgttcgagtggctacgccacagaacaaccggacgccctgcggtgtgg
optimized_alkS-CR          accgtggctcttgcgttcgagtggctacgccacagaacaaccggacgccctgcggtgtgg
                           *********************************************** ***** ******
                                                                      AatII  PstI

                           IleSerLeuArgAlaSerSerTyrSerGluPheAspIleCysAlaGluIleIleGluGln
native_alkS                atttctttaagagccagttcttacagtgaatttgatatctgcgcagaaattattgagcag
optimized_alkS-ED          atttctttaagagccagttcttacagtgaatttgatatctgcgcagaaattattgagcag
optimized_alkS-CR          atttctttaagagccagttcttacagtgaatttgatatctgcgcagaaattattgagcag
                           ************************************************************

                           LeuGluAlaPheGluLeuValThrPheSerHisValArgGluGlyValSerLysProThr
native_alkS                cttgaagcgttcgaactggtaacattcagccatgtgagagagggtgtgagcaagcctacg
optimized_alkS-ED          cttgaagcgttcgaactggtaacattcagccatgtgagagagggtgtgagcaagcctacg
optimized_alkS-CR          cttgaagcgttcgaactggtaacattcagccatgtgagagagggtgtgagcaagcctacg
                           ************************************************************

                           LeuLeuArgAspLeuAlaSerSerLeuTrpGlnSerThrSerSerAsnGluIleGluThr
native_alkS                ctcttgcgagaccttgcatccagtctttggcagagtacctcgagtaacgaaatagaaacg
optimized_alkS-ED          ctcttgcgagaccttgcatccagtctttggcagagtacctcgagtaacgaaatagaaacg
optimized_alkS-CR          ctcttgcgagaccttgcatccagtctttggcagagtacctcgagtaacgaaatagaaacg
                           ************************************************************

                           LeuIleCysLeuAspAsnIleAsnGlnGlyLeuGlyLeuProLeuLeuHisAlaLeuMet
native_alkS                ctaatttgtttggataatattaatcagggcttgggcttgccgttgttgcacgcgcttatg
optimized_alkS-ED          ctaatttgtttggataatattaatcagggcttgggcttgccgttgttgcacgcgcttatg
optimized_alkS-CR          ctaatttgtttggataatattaatcagggcttgggcttgccgttgttgcacgcgcttatg
                           ************************************************************

                           GluPheMetLeuGluThrProLysSerIleArgPheAlaValAlaGlyAsnThrIleLys
native_alkS                gagttcatgttagaaacaccaaaaagtatcaggtttgcagtcgcaggtaatacaataaaa
optimized_alkS-ED          gagttcatgttagaaacaccaaaaagtatcaggtttgcagtcgcaggtaatacaataaaa
optimized_alkS-CR          gagttcatgttagaaacaccaaaaagtatcaggtttgcagtcgcaggtaatacaataaaa
                           ************************************************************

                           GlyPheSerArgLeuLysLeuAlaGlyAlaMetGlnGluHisThrGluLysAspLeuAla
```

```
native_alkS                 gggttctcgcggcttaaacttgcaggcgctatgcaggagcacaccgagaaagatttggcc
optimized_alkS-ED           gggttctcgcggcttaaacttgcaggcgctatgcaggagcacaccgagaaagatttggcc
optimized_alkS-CR           gggttctcgcggcttaaacttgcaggcgctatgcaggagcacaccgagaaagatttggcc
                            ************************************************************

                            PheSerAlaAspGluAlaValAlaLeuValGluAlaGluAlaValLeuGlyValSerGlu
native_alkS                 tttagcgcagacgaggcagtggcgttagtggaggcagaggctgttcttggagtttctgag
optimized_alkS-ED           tttagcgcagacgaggcagtggcgttagtggaggcagaggctgttcttggagtttctgag
optimized_alkS-CR           tttagcgcagacgaggcagtggcgttagtggaggcagaggctgttcttggagtttctgag
                            ************************************************************
                            ValGlnIleGluAlaLeuValGlnGluMetGluGlyTrpProValLeuIleGlyPheLeu
native_alkS                 gtacagatagaggccttggtgcaagaaatggagggggtggcctgttcttatcgggtttttg
optimized_alkS-ED           gtacagatagaggccttggtgcaagaaatggagggggtggcctgttcttatcgggtttttg
optimized_alkS-CR           gtacagatagaggccttggtgcaagaaatggagggggtggcctgttcttatcgggtttttg
                            ************************************************************

                            LeuLysCysGluLeuProAlaLysSerIleSerThrValValGluIleAspAsnTyrPhe
native_alkS                 ttaaaatgtgagttgccggccaagtctatttcaacagtagttgaaatagacaattacttt
optimized_alkS-ED           ttaaaatgtgagttgccggccaagtctatttcaacagtagttgaaatagacaattacttt
optimized_alkS-CR           ttaaaatgtgagttgccggccaagtctatttcaacagtagttgaaatagacaattacttt
                            ************************************************************

                            AsnAspGluIlePheGluAlaLeuProGluArgTyrArgValPheLeuValAsnSerSer
native_alkS                 aatgatgaaatatttgaggcgcttcccgagcggtatcgtgtttttcttgtaaattcttca
optimized_alkS-ED           aatgatgaaatatttgaggcgcttcccgagcggtatcgtgtttttcttgtaaattcttca
optimized_alkS-CR           aatgatgaaatatttgaggcgcttcccgagcggtatcgtgtttttcttgtaaattcttca
                            ************************************************************

                            LeuLeuAspValValThrProAspGlyTyrAsnTyrValPheLysCysValAsnAlaAla
native_alkS                 ttgctcgatgtcgtgacgcctgatggctacaattatgtattcaaatgcgtcaatgcggcc
optimized_alkS-ED           ttgctcgatgtcgtgacgcctgatggctacaattatgtattcaaatgcgtcaatgcggcc
optimized_alkS-CR           ttgctcgatgtcgtgacgcctgatggctacaattatgtattcaaatgcgtcaatgcggcc
                            ************************************************************

                            SerCysIleLysTyrLeuSerThrAsnTyrMetLeuLeuArgHisValGluGlyGluPro
native_alkS                 tcatgtattaaatatctgagcacgaattacatgttacttcgccatgtagaaggtgagccg
optimized_alkS-ED           tcatgtattaaatatctgagcacgaattacatgttacttcgccatgtagaaggtgagccg
optimized_alkS-CR           tcatgtattaaatatctgagcacgaattacatgttacttcgccatgtagaaggtgagccg
                            ************************************************************

                            AlaGlnPheThrLeuHisProValLeuArgAspPheLeuGlnGlyIleAlaTrpAlaGlu
native_alkS                 gcgcagtttacactacatccagtattgcgcgattttcttcaaggaattgcttgggctgaa
optimized_alkS-ED           gcgcagtttacactacatccagtattgcgcgattttcttcaaggaattgcttgggctgaa
optimized_alkS-CR           gcgcagtttacactacatccagtattgcgcgattttcttcaaggaattgcttgggctgaa
                            ************************************************************

                            AsnProAlaLysArgSerTyrLeuLeuLysArgAlaAlaPheTrpHisTrpArgArgGly
native_alkS                 aatcctgctaaaagatcctacttgcttaagcgcgcagctttctggcattggcgtagaggc
optimized_alkS-ED           aatcctgctaaaagatcctacttgcttaagcgcgcagctttctggcattggcgtagaggc
optimized_alkS-CR           aatcctgctaaaagatcctacttgcttaagcgcgcagctttctggcattggcgtagaggc
                            ************************************************************
                            GluTyrGlnTyrAlaIleArgIleAlaLeuArgAlaAsnAspCysArgTrpAlaValGly
native_alkS                 gaataccagtacgcaatacgaatagccctacgggcgaatgactgtcgctgggctgtcggc
optimized_alkS-ED           gaataccagtacgcaatacgaatagccctacgggcgaatgactgtcgctgggctgtcggc
optimized_alkS-CR           gaataccagtacgcaatacgaatagccctacgggcgaatgactgtcgctgggctgtcggc
                            ************************************************************

                            MetSerGluGlyIleIleLeuAspLeuSerPheArgGlnGlyGluIleAspThrLeuArg
native_alkS                 atgagtgagggaataattttagatttgtcatttcgtcagggcgaaatagatacgctgaga
optimized_alkS-ED           atgagtgagggaataattttagatttgtcatttcgtcagggcgaaatagatacgctgaga
optimized_alkS-CR           atgagtgagggaataattttagatttgtcatttcgtcagggcgaaatagatacgctgaga
                            ************************************************************

                            HisTrpLeuSerGluLeuProValLysAspLeuHisLysAsnProIleValLeuIleCys
native_alkS                 cactggctgtcggagctgccagtgaaggacttgcacaagaaccccatagtacttatttgt
optimized_alkS-ED           cactggctgtcggagctgccagtgaaggacttgcacaagaaccccatagtacttatttgt
optimized_alkS-CR           cactggctgtcggagctgccagtgaaggacttgcacaagaaccccatagtacttatttgt
```

```
                        ************************************************************

                        PheAlaTrpValLeuTyrPheSerGlnGlnSerAlaArgAlaGluLysLeuLeuLysAsp
native_alkS             ttcgcgtgggtattgtatttcagtcagcaaagcgcgcgagcagagaagttacttaaagac
optimized_alkS-ED       ttcgcgtgggtattgtatttcagtcagcaaagcgcgcgagcagagaagttacttaaagac
optimized_alkS-CR       ttcgcgtgggtattgtatttcagtcagcaaagcgcgcgagcagagaagttacttaaagac
                        ************************************************************

                        LeuIleThrProProAspLysLysAsnLysTrpGlnGluLysGlyTrpProGlnLeuVal
native_alkS             ctaattaccccgcccgataaaaaaaacaaatggcaagaaaaaggatggccgcagcttgtg
optimized_alkS-ED       ctaattaccccgcccgataaaaaaaacaaatggcaagaaaaaggatggccgcagcttgtg
optimized_alkS-CR       ctaattaccccgcccgataaaaaaaacaaatggcaagaaaaaggatggccgcagcttgtg
                        ************************************************************

                        PheAlaIleGlyLysAlaThrAsnAspGluMetLeuLeuSerGluGluLeuCysAsnLys
native_alkS             tttgcaataggtaaagcaacgaatgatgaaatgcttttgagtgaggagctctgtaataag
optimized_alkS-ED       tttgcaataggtaaagcaacgaatgatgaaatgcttttgagtgaggagctgtgtaataag
optimized_alkS-CR       tttgcaataggtaaagcaacgaatgatgaaatgcttttgagtgaggagctgtgtaataag
                        ************************************************** *********
                                                                      SacI

                        TrpIleSerLeuPheGlyAspSerAsnAlaValGlyLysGlyAlaAlaLeuThrCysLeu
native_alkS             tggattagtttatttggggattcaaacgcagttgggaaaggggccgcgctaacctgcttg
optimized_alkS-ED       tggattagtttatttggggattcaaacgcagttgggaaaggggccgcgctaacctgcttg
optimized_alkS-CR       tggattagtttatttggggattcaaacgcagttgggaaaggggccgcgctaacctgcttg
                        ************************************************************

                        AlaPheIlePheAlaSerGluTyrArgPheAlaGluLeuGluLysValLeuAlaGlnAla
native_alkS             gcttttatttttgccagtgagtatagatttgcagagttagagaaggtgctggctcaggcc
optimized_alkS-ED       gcttttatttttgccagtgagtatagatttgcagagttagagaaggtgctggctcaggcc
optimized_alkS-CR       gcttttatttttgccagtgagtatagatttgcagagttagagaaggtgctggctcaggcc
                        ************************************************************

                        GlnAlaValAsnLysPheAlaLysGlnAspPheAlaPheGlyTrpLeuTyrValAlaLys
native_alkS             caagccgtgaataaatttgcaaaacaagattttgctttcggttggctgtatgtcgccaag
optimized_alkS-ED       caagccgtgaataaatttgcaaaacaagattttgctttcggttggctgtatgtcgccaag
optimized_alkS-CR       caagccgtgaataaatttgcaaaacaagattttgctttcggttggctgtatgtcgccaag
                        ************************************************************

                        LeuGlnGlnAlaLeuAlaSerGlyLysMetSerTrpAlaArgGlnLeuIleThrGlnAla
native_alkS             cttcaacaagcgctagcaagcgggaaaatgagctgggccaggcagcttataactcaagca
optimized_alkS-ED       ctgcaacaagcgctagcaagcgggaaaatgagctgggccaggcagcttataactcaagca
optimized_alkS-CR       ctgcaacaagcgctagcaagcgggaaaatgagctgggccaggcagcttataactcaagca
                        ** *********************************************************
                        HindIII

                        ArgThrAspSerGlyAlaGlnIleMetGluThrAlaPheThrSerLysMetLeuAspAla
native_alkS             cgcacagacagtggcgcgcagattatggagaccgcgtttacctcgaaaatgctggacgct
optimized_alkS-ED       cgcacagacagtggcgcgcagattatggagaccgcgtttacctcgaaaatgctggacgct
optimized_alkS-CR       cgcacagacagtggcgcgcagattatggagaccgcgtttacctcgaaaatgctggacgct
                        ************************************************************

                        LeuGluLeuGluSerAsnTyrGluLeuCysArgLeuAspThrSerGluGluLysPheSer
native_alkS             ctagagcttgagtcaaattatgaattgtgccgcttggacacctcagaagaaaagttctcc
optimized_alkS-ED       ctggagcttgagtcaaattatgaattgtgccgcttggacacctcagaagaaaagttctcc
optimized_alkS-CR       ctggagcttgagtcaaattatgaattgtgccgcttggacacctcagaagaaaagttctcc
                        ** *********************************************************
                        XbaI

                        GluIleLeuGluPheIleAlaAsnHisGlyValThrAspValPhePheSerValCysArg
native_alkS             gaaattttagagtttattgccaatcacggggtgacagacgtgtttttttccgtatgccgt
optimized_alkS-ED       gaaattttagagtttattgccaatcacggggtgacagacgtgtttttttccgtatgccgt
optimized_alkS-CR       gaaattttagagtttattgccaatcacggggtgacagacgtgtttttttccgtatgccgt
                        ************************************************************

                        ValValSerAlaTrpArgLeuGlyArgAsnAspLeuAsnGlySerIleGluIleLeuGlu
native_alkS             gttgtgtcagcttggcggcttggaaggaatgacctgaatggttccattgagattttggag
optimized_alkS-ED       gttgtgtcagcttggcggcttggaaggaatgacctgaatggttccattgagattttggag
optimized_alkS-CR       gttgtgtcagcttggcggcttggaaggaatgacctgaatggttccattgagattttggag
```

```
                     ************************************************************

                     TrpAlaLysAlaTyrAlaAlaGluLysAsnLeuProArgLeuGluValMetSerGlnIle
native_alkS          tgggcgaaggcatatgcggctgaaaaaaatctaccaagattggaagttatgagccaaatt
optimized_alkS-ED    tgggcgaaggcatatgcggctgaaaaaaatctaccaagattggaagttatgagccaaatt
optimized_alkS-CR    tgggcgaaggcatatgcggctgaaaaaaatctaccaagattggaagttatgagccaaatt
                     ************************************************************

                     GluIleTyrGlnArgLeuLeuPheGlnGlyValThrTyrIleAsnThrLeuGlnAlaPhe
native_alkS          gagatctatcaacgcttgctctttcaaggcgtaacgtacataaatacgttacaagctttt
optimized_alkS-ED    gagatctatcaacgcttgctctttcaaggcgtaacgtacataaatacgttacaagccttt
optimized_alkS-CR    gagatctatcaacgcttgctctttcaaggcgtaacgtacataaatacgttacaagccttt
                     ******************************************************** ***
                                                                        HindIII

                     GluAspArgLysIlePheSerGlyProHisSerAlaProLeuLysAlaArgLeuLeuLeu
native_alkS          gaagatcgcaagattttctccggaccgcactcagcccccctaaaggcacgcctgctgctt
optimized_alkS-ED    gaagatcgcaagattttctccggaccgcactcagcccccctaaaggcacgcctgctgctt
optimized_alkS-CR    gaagatcgcaagattttctccggaccgcactcagcccccctaaaggcacgcctgctgctt
                     ************************************************************

                     ValGlnSerLeuAlaLeuSerArgAspGlnAsnPheHisLeuAlaAlaHisArgAlaLeu
native_alkS          gttcaatcactcgcgctttcccgagatcagaactttcatcttgccgcgcacagagcgcta
optimized_alkS-ED    gttcaatcactcgcgctttcccgagatcagaactttcatcttgccgcgcacagagcgcta
optimized_alkS-CR    gttcaatcactcgcgctttcccgagatcagaactttcatcttgccgcgcacagagcgcta
                     ************************************************************

                     LeuAlaIleGlnGlnAlaArgLysIleSerAlaGlyGlnLeuGluValArgGlyLeuLeu
native_alkS          ttggctattcagcaagcccgtaaaattagcgcaggccaattggaagtccgtggcttattg
optimized_alkS-ED    ttggctattcagcaagcccgtaaaattagcgcaggccaattggaagtccgtggcttattg
optimized_alkS-CR    ttggctattcagcaagcccgtaaaattagcgcaggccaattggaagtccgtggcttattg
                     ************************************************************

                     TyrLeuAlaGlyAlaGlnAlaGlyGlyGlyThrLeuLysLysAlaGlnHisAsnIleAla
native_alkS          tatttagccggagcgcaagcaggtggcggcacattaaagaaggctcagcataacattgct
optimized_alkS-ED    tatttagccggagcgcaagcaggtggcggcacattaaagaaggctcagcataacattgct
optimized_alkS-CR    tatttagccggagcgcaagcaggtggcggcacattaaagaaggctcagcataacattgct
                     ************************************************************

                     TyrAlaLeuGluMetAlaLysGlnLeuGlnCysPheGlnThrValLeuAspGluIleArg
native_alkS          tatgcattggagatggcaaaacagcttcaatgctttcaaacagtgctggatgaaatacgt
optimized_alkS-ED    tatgcattggagatggcaaaacagcttcaatgctttcaaacagtgctggatgaaatacgt
optimized_alkS-CR    tatgcattggagatggcaaaacagcttcaatgctttcaaacagtgctggatgaaatacgt
                     ************************************************************

                     LeuIleLysArgLeuIleProThrSerCysGluValPheAlaAlaValAsnLeuAspGln
native_alkS          ttgattaagagattaataccgacttcatgtgaagtcttcgcagcagttaatttagatcaa
optimized_alkS-ED    ttgattaagagattaataccgacttcatgtgaagtcttcgcagcagttaatttagatcaa
optimized_alkS-CR    ttgattaagagattaataccgacttcatgtgaagtcttcgcagcagttaatttagatcaa
                     ************************************************************

                     AlaIleGlyAlaPheSerLeuProGlnMetValGluIleArgLysSerAlaGluAsnLys
native_alkS          gcgattggagcttttagtctgccgcaaatggtggagattagaaaatccgcagaaaataaa
optimized_alkS-ED    gcgattggagcttttagtctgccgcaaatggtggagattagaaaatccgcagaaaataaa
optimized_alkS-CR    gcgattggagcttttagtctgccgcaaatggtggagattagaaaatccgcagaaaataaa
                     ************************************************************

                     AlaGlyAspPheLeuThrLeuLysGlnValSerValLeuLysLeuValLysGluGlyCys
native_alkS          gctggtgattttttgacactgaagcaggtcagtgtcttgaaacttgtaaaagaggggtgc
optimized_alkS-ED    gctggtgattttttgacactgaagcaggtcagtgtcttgaaacttgtaaaagaggggtgc
optimized_alkS-CR    gctggtgattttttgacactgaagcaggtcagtgtcttgaaacttgtaaaagaggggtgc
                     ************************************************************

                     SerAsnLysGlnIleAlaThrLysMetTyrValThrGluAspAlaIleLysTrpHisMet
native_alkS          tcaaacaaacaaatagcaacaaagatgtatgtcaccgaagatgctataaagtggcatatg
optimized_alkS-ED    tcaaacaaacaaatagcaacaaagatgtatgtcaccgaagatgctataaagtggcatatg
```

```
optimized_alkS-CR       tcaaacaaacaaatagcaacaaagatgtatgtcaccgaagatgctataaagtggcatatg
                        ************************************************************


                        ArgLysIlePheThrIleLeuAsnValLysSerArgThrGlnAlaIleIleGluAlaGlu
native_alkS             aggaaaatatttaccatccttaatgtaaagagtcgcacgcaagcaataattgaagccgaa
optimized_alkS-ED       aggaaaatatttaccatccttaatgtaaagagtcgcacgcaagcaataattgaagccgaa
optimized_alkS-CR       aggaaaatatttaccatccttaatgtaaagagtcgcacgcaagcaataattgaagccgaa
                        ************************************************************


                        ArgGlnGlyValIleEnd
native_alkS             cgtcagggtgttatctga
optimized_alkS-ED       cgtcagggtgttatctga
optimized_alkS-CR       cgtcagggtgttatctga
                        ******************
```

Note removal of restriction sites incompatible with the SEVA standard (red), changes to eliminate the Crc site (yellow) and optimization of some codons throughout the sequence.

**Supplementary Figure S2.** Organization of AlkS/PalkB-based expression vectors

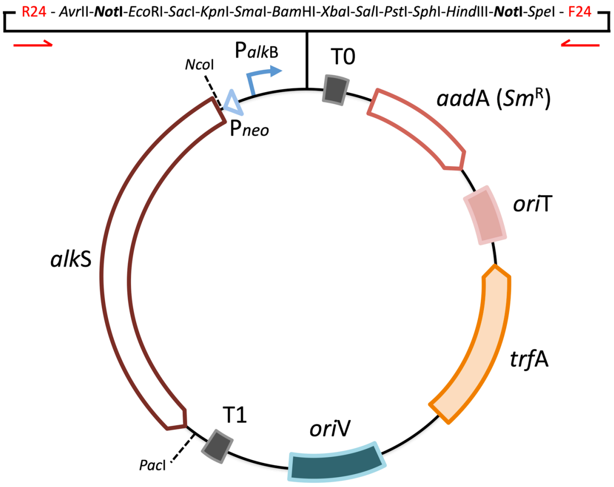


*alkS*$^{ED}$ : pSEVA429 *crc*$^{+}$

*alkS*$^{CR}$ : pSEVA429

The drawing shows the arrangement of the expression device in the frame of a SEVA-formatted plasmid. Depending on whether the edited sequence of the regulator keeps (*alkS*$^{ED}$) or not (*alkS*$^{CR}$) the Crc-binding site in its mRNA the name of the standardized plasmids changes as indicated.

**Supplementary Figure S3.** Functional insert of plasmids pJAMA30 and pAR*alkS*

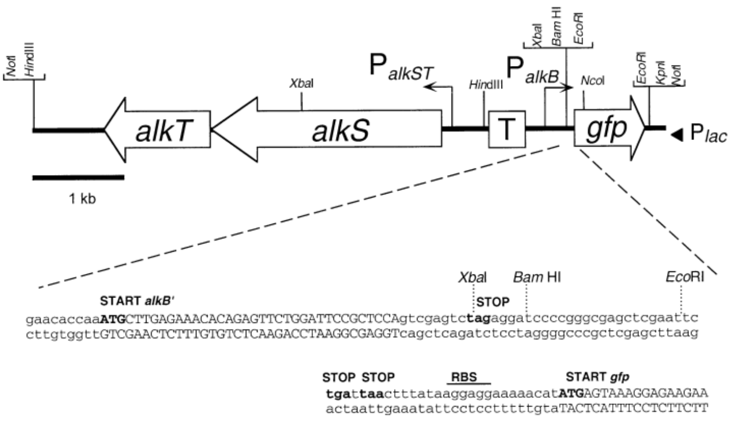


The expression cassette of the cargo borne by plasmid pJAMA30 is shown with an indication of all functional parts. The same DNA segment was cloned in a SEVA vector as NotI fragment to produce pAR*alkS* (see main text for explanation). Figure reproduced from (1).
